# Zika virus infection at mid-gestation results in fetal cerebral cortical injury and fetal death in the olive baboon

**DOI:** 10.1101/331108

**Authors:** Sunam Gurung, Nicole Reuter, Alisha Preno, Jamie Dubaut, Hugh Nadeau, Kimberly Hyatt, Krista Singleton, Ashley Martin, W. Tony Parks, James F. Papin, Dean A. Myers

**Affiliations:** Department of Obstetrics and Gynecology, University of Oklahoma Health Sciences Center, Oklahoma City, OK 73109; Division of Comparative Medicine, University of Oklahoma Health Sciences Center, Oklahoma City, OK 73109; Department of Physiology, University of Oklahoma Health Sciences Center, Oklahoma City, OK 73109; Department of Pathology, Northwestern University, Chicago, IL

## Abstract

Zika virus (ZIKV) infection during pregnancy in humans is associated with an increased incidence of congenital anomalies including microcephaly as well as fetal death and miscarriage and collectively has been referred to a Congenital Zika Syndrome (CZS). Animal models for ZIKV infection in pregnancy have been developed including mice and macaques. While microcephaly has been achieved in mice via direct injection of ZIKV into the fetal brain or via interference with interferon signaling, in macaques the primary fetal CZS outcome are ocular defects. In the present study we develope the olive baboon (*Papio anubis*), as a model for the vertical transfer of ZIKV during pregnancy. We infected four mid-gestation, timed-pregnant baboons with the French Polynesian ZIKV isolate (10^4^ ffu) and examined the acute phase of vertical transfer by stopping the study of one dam at 7 days post infection (dpi), two at 14 dpi and one at 21 dpi. All dams exhibited mild to moderate rash and conjunctivitis; three of four dams exhibited viremia at 7 dpi. Of the three dams studied to 14 to 21 days, only one still exhibited viremia on day 14. Vertical transfer of ZIKV to the fetus was found in two pregnancies; in one, vertical transfer was associated with fetal death at ∼14 dpi. In the other, vertical transfer was observed at 21 dpi. Both fetuses had ZIKV RNA in the fetal cerebral cortex as well as other tissues. The 21 dpi fetal cerebral cortex exhibited notable defects in radial glia, radial glial fibers, loss and or damage of immature oligodendrocytes and a loss in neuroprogenitor cells (NPCs). In addition, indices of pronounced neuroinflammation were observed including astrogliosis, increased microglia and IL-6 expression. The dams studied to 14 dpi (n=2) and 21 dpi (n=1) exhibited a anti-ZIKV IgM response and IgG response (21 dpi) that included transfer of the IgG to the fetal compartment (cord blood). The severity of systemic inflammatory response (cytokines and chemokines) reflected the vertical transfer of ZIKV in the two pregnancies. As such, these events likely represent the early mechanisms that lead to microcephaly and/or other CNS pathologies in a primate infected with ZIKV and are the first to be described in a non-human primate during the acute phase of ZIKV infection with a contemporaneous ZIKV strain. The baboon thus represents a major NHP for advancing as a model for ZIKV induced brain pathologies to contrast and compare to humans as well as other NHPs such as macaques.

**AUTHOR SUMMARY:** Zika virus is endemic in the Americas, primarily spread through mosquitos and sexual contact. Zika virus infection during pregnancy in women is associated with a variety of fetal pathologies now referred to as Congenital Zika Syndrome (CZS), with the most severe pathology being fetal microcephaly. Developing model organisms that faithfully recreate Zika infection in humans is critical for future development of treatments and preventions. In our present study, we infected Olive baboons at mid-gestation with Zika virus and studied the acute period of viremia and transfer of Zika virus to the fetus during the first three weeks after infection to better understand the timing and mechanisms leading to CZS. We observed Zika virus transfer to fetuses resulting in fetal death in one pregnancy and in a second pregnancy, significant damage to the frontal cortex of the fetal brain consistent with development of microcephaly, closely resembling infection in pregnant women. Our baboon model differs from macaque non-human primate models where the primary fetal outcome during pregnancy following infection with contemporary strains of Zika virus is ocular pathology. Thus, the baboon provides a promising new non-human primate model to further compare and contrast the consequences of Zika virus infection in pregnancy to humans and macaques to better understand the disease.

## INTRODUCTION

Originally isolated from a febrile sentinel rhesus monkey in the Zika forest in Uganda in 1947, Zika virus (ZIKV) belongs to the *Flaviviridae* family, genus Flavivirus, which includes dengue (DENV), West Nile (WNV), yellow fever (YFV), and Japanese encephalitis virus (JEV) (1, 2). Prior to the 2007 ZIKV outbreak on Yap Island (Federated States of Micronesia; ∼5000 cases), there had been only 14 known cases of ZIKV infection, restricted to Africa and Southeast Asia. These early cases of ZIKV infection, including the Yap Island outbreak, were largely self-limiting, relatively harmless with symptoms of mild fever, rash, conjunctivitis, arthritis and arthralgia (3). A subsequent outbreak of ZIKV in French Polynesia (2013-14) resulting in ∼30,000 cases (∼11% of the population) was associated with a decided shift in symptomology including an increased incidence of Guillain-Barre syndrome (GBS) (4). The ensuing ZIKV epidemic in Brazil (2014-15) that spread through parts of South and Central America was larger in scope and alarmingly linked to an increased incidence of newborns with microcephaly and a variety of other congenital anomalies (5-8). A retrospective analysis of the French Polynesian outbreak confirmed the association between ZIKV and increased incidence of microcephaly (5, 9). The first documented cases of intrauterine (vertical) transmission of ZIKV were reported in 2016 in Brazil (8). In addition to a spectrum of congenital malformations, now collectively referred to as Congenital Zika Syndrome (CZS), ZIKV infection in pregnancy is associated with intrauterine fetal demise and increased incidence of miscarriage and preterm birth (5, 6).

The development of animal models that faithfully recapitulate the complex pathogenesis of ZIKV infection that leads to trans-placental passage of the virus and fetal CNS damage and accompanying CZS anomalies is essential for developing and testing vaccines and anti-viral strategies. Studies of ZIKV infection in mouse models have largely relied upon mice deficient in interferon (IFN) signaling; either transgenic mice lacking the ability to produce or respond to IFNs(10-12) or wild-type mice passively immunized against Type 1 IFNs (11). In pregnant mice lacking type 1 IFN signaling (Ifnar1^−/-^) or following antibody neutralization of Ifnar-1 signaling, infection with the French Polynesian clinical isolate (H/PF/2013) resulted in placental and fetal infection leading to a range of fetal pathologies including fetal death and/or intrauterine growth restriction (12). ZIKV infection has also been achieved in wild-type mice following direct intrauterine (intracranial) infection of the embryonic pups with either Asian or Mexican origin ZIKV (13, 14). These studies reported ZIKV targeting of neuroprogenitor cells (NPCs), radial glia and immature neurons, as well as neuroinflammation (microglia activation) resulting in cortical damage in the fetus resembling human microcephaly. ZIKV infection of neonatal wild-type mice has also been accomplished either via direct intracranial injection of virus (15), or peripheral infection (16) using various strains of ZIKV including African, Asian and American strains leading to a spectrum of CNS pathologies. Fetal infection has also been reported in wild-type mice after delivery of the virus directly into the uterine wall (17). This study reported fetal infection and reduced cerebral cortical thickness in the pups when the dams were infected at embryonic day 10 but not 14, suggesting that a gain in placental IFN signaling, as has been reported in term gestation human placentas, may provide a barrier to ZIKV (18, 19). The necessity of circumventing IFN signaling in pregnant mice in order to achieve vertical transmission of ZIKV limits the translational application to human pregnancy. Curiously, even ancestral strains of ZIKV (African) not associated with adverse fetal outcome in humans and that are usually asymptomatic or mildly symptomatic in adults, induce fetal CNS damage in these mice.

Non-human primates (NHPs) are the best documented animal reservoirs for ZIKV (and related flaviviruses). ZIKV infection has been achieved in male, and non-pregnant and pregnant female rhesus (*Macacca mulatta*)(20-31) cynomolgus (*Macacca fascicularis*) (22, and pigtail macaques (*Macacca nemestrina*) (33) following subcutaneous (SC) inoculation with French Polynesian (H/PF/2013),(20) Puerto Rican (PRVABC59, 2015) (21, 22, 30-32), Brazilian (Brazil/ZKV2015; Zika virus/H.sapiens-tc/BRA/2015/Brazil_SPH2015) (26-29) and Cambodian strains of ZIKV (FSS13025) (32). There are now several reports of ZIKV infection in pregnant macaques, with one study describing infection of a single pigtail macaque using an Asian strain (FSS13025 strain; Cambodia, 2010) (33), and four studies with pregnant rhesus macaques infected with either the French Polynesian or Puerto Rican strains (20, 23, 30, 31). In the study of a single pigtail macaque infected at the gestational equivalent to human early 3^rd^ trimester (33), a significant fetal pathological CNS outcome at near-term gestation (43 days post-infection), albeit the dose of ZIKV used was substantially greater (five SC inoculations, 1×10^7^ pfu/site) than expected from a mosquito bite (∼1×10^3^ to 1×10^4^ pfu) (34). In this study, fetal brain malformations/lesions primarily in white matter (periventricular lesions, cerebral white matter hypoplasia and gliosis), with normal cortical folding and no evidence of cortical malformations were noted at term gestation. The fetal CNS pathology was observed despite using a ZIKV strain not associated with adverse pregnancy outcome, fetal microcephaly or CZS in humans. A study of pregnant rhesus macaques infected with the French Polynesian strain of ZIKV (1×10^4^ pfu) during either mid-1^st^ or early 3^rd^ trimesters (two dams each) resulted in 100% vertical transfer of the virus to the fetus (35). This study noted normal fetal cerebral cortical volume, thickness and sulcation with no histopathological evidence of inflammation. At the termination of the study, ZIKV RNA was detected in various fetal tissues but not brain. These investigators did note ocular pathologies similar to that described in human fetuses from ZIKV infected mothers as well as evidence of inflammation at the maternal-fetal placental interface. A recent study describing infection of five rhesus macaques with the Puerto Rican strain during either the 1^st^ or 2^nd^ trimesters reported uterine vasculitis, placental villous damage, and evidence of abnormal oxygen transport within the placenta by the 3^rd^ trimester (31). Subtle effects on fetal brain structure were noted in two of the five pregnancies as observed by MRI analysis in the third trimester (135 dGA), and ZIKV RNA was detected in the fetal cerebral cortex. A recent study of a single rhesus macaque infected with the Puerto Rican strain during the 1^st^ trimester resulted in fetal demise by 49 dpi, again consistent with reports of fetal death in human pregnancies infected with the virus (30). This study noted presence of ZIKV RNA in fetal tissues, including low copy numbers in the fetal brain, yet no apparent fetal CNS anomalies. Overall, the macaque species appear highly permissive to ZIKV infection with no apparent barrier to vertical transfer of ZIKV across the placenta at any stage of pregnancy examined to date. The effects on ZIKV on the fetal CNS from these studies are equivocal, ranging from severe to none, and could be related to viral dosing or macaque subspecies. Despite these elegant and essential studies describing ZIKV infection in pregnant macaques, there is still a clear need to develop additional NHP models to study pregnancy and fetal outcome from ZIKV infection to further explore the timing of gestational exposure to the virus and maternal-fetal responses to the viral infections.

In the present study, we developed the olive baboon (*Papio anubis*) as an alternative NHP model to study ZIKV infection and pathogenicity during pregnancy that can be correlated with the human situation and compared/contrasted with ZIKV infection in pregnant macaques. Our focus was on the acute period following ZIKV infection (first three weeks post-infection) and early events in transplacental passage and pathogenesis of the fetal brain. The olive baboon is similar to humans in terms of size, genetics, reproduction, brain development and immune repertoire makes the baboon an excellent translational model to study ZIKV infection and for vaccine and therapeutics development (36-38). The baboon has been used as a NHP model for assessing safety and efficacy of vaccines in adults, pregnant females and their infants (37, 39). The baboon is permissive to flavivirus infection and replication and produces a virus-specific immune response (38, 40). Herein, we describe infection of four timed-pregnant olive baboons at mid-gestation with a contemporary French Polynesian strain of ZIKV (H/PF/2013). The French Polynesian ZIKV strain contains a single point mutation in the prM protein that dramatically increases ZIKV infectivity in both human and mouse neuroprogenitor cells compared to the ancestral African/Asian ZIKV strains(41). This mutation was conserved during the ZIKV spread through the Americas and is associated with adverse fetal outcomes (42-44). We report that the pregnant olive baboon is susceptible to ZIKV infection during gestation including vertical transfer of virus to the fetus resulting in both fetal death as well as fetal cerebral cortical pathologies consistent with potential for development of microcephaly.

## RESULTS

- **Symptomology and pregnancy outcome.**

All ZIKV infected dams had minor weight loss during the study period (Dam 1: 16.4 start, 16.2 kg end [7 days]; Dam 2 16.0 start, 15.6 kg end [14 days]; Dam 3: 21 start, 20.9 kg end; 14 days; Dam 4: 13.8 start, 13.6 kg end [21 days]), however, none of the dams exhibited inappetence. Dam 1 exhibited a mild rash on day three post-infection in the axillary and inguinal regions as well as conjunctivitis that cleared by day seven post-infection; Dam 2 exhibited a mild rash in the axillary and inguinal regions and minor conjunctivitis by day three post-infection which expanded to moderate to severe maculopapular rash on the abdomen and inguinal regions with mild rash on the chest and back of both arms with mild conjunctivitis by day seven that resolved by day 14 post-infection. Dam 3 developed a mild rash in the axillary region and a moderate rash in the inguinal region with moderate conjunctivitis that resolved by day seven post-infection. Dam 4 exhibited a mild rash on day three post-infection in the axillary and inguinal regions as well as mild conjunctivitis that progressed to a mild to moderate rash by day seven-post infection that included the abdomen, chest and backs of arms that resolved by 14 days post-infection. Body temperatures obtained under ketamine sedation did not show any fever greater than 1°C above day 0 over the course of the study for each animal.

Complete blood counts (CBCs) were evaluated for all females on EDTA-anticoagulated whole blood samples collected on day 0 and subsequent days-post infection as shown in the experimental timeline (Idexx ProCyte DX hematology analyzer; Idexx laboratories, ME). CBC’s included analysis for red blood cells (RBCs), hemoglobin, hematocrit and platelet count. RBC, hemoglobin and hematocrit numbers did not show any differences pre-and post ZIKV infection for any of the infected females. Platelet counts did not change in response to ZIKV infection in any dam.

Normal fetal heart rates were obtained for Dams 1, 2 and 4 from the day of inoculation through pregnancy termination on days 7, 14 and 21 respectively. Dam 3 exhibited normal fetal heart rate through day 7 post-infection. However, on day 14, no fetal heart rate was found and at subsequent necropsy, this fetus was still born with fetal demise likely within the preceding 24 hours. There was rupture of the fetal membranes in this pregnancy as well and signs of meconium staining.

- **Maternal ZIKV Viral loads**

For Dams 1, 2 and 4, ZIKV RNA was not detected in whole blood on day three but was detected on day seven post-infection with resolution by day 14 post-infection (Fig 2A). In Dam 3, ZIKV RNA was detected on days 3 through 14 post-infection (study termination, Fig 2A). Peak viremia in the four dams ranged from 8.5 ×; 10^3^ through 2.2 ×; 10^6^ copies/ml blood. ZIKV RNA was detected in saliva from Dams 1 and 4 at seven days post-infection, and from Dam 2 on day 14 post-infection (Fig 2B).

**Figure 1.**
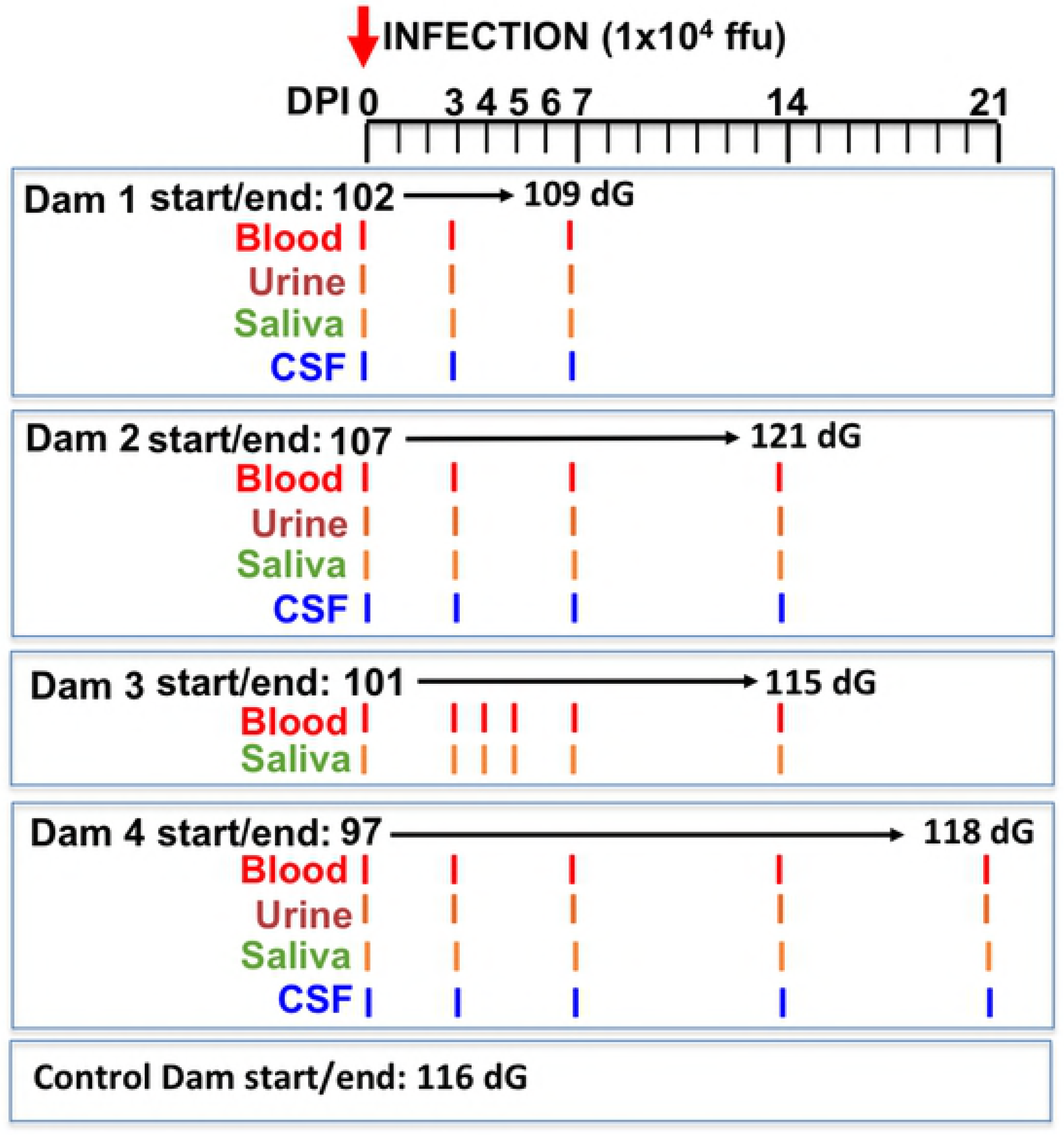
Schematic representation of the experimental design. Multiparous timed pregnant olive baboons (n=4) were infected subcutaneously (1×10^4^ ffu, 1 ml volume, strain H/FP/2103) on Day 0. Maternal blood, urine, CSF and saliva were obtained on the indicated days for each animal. The gestational age at the time of infection ranged from 97-107 days gestation (dG). One dam was euthanized at day 7 post infection (Dam 1), two dams on day 14 post infection (Dams 2,3) and on at 21 days post infection for collection of maternal and fetal tissues. The gestational age range at necropsy was 109-121 dG; a timed pregnant control dam was euthanized at 116 dG.

**Figure 2.**
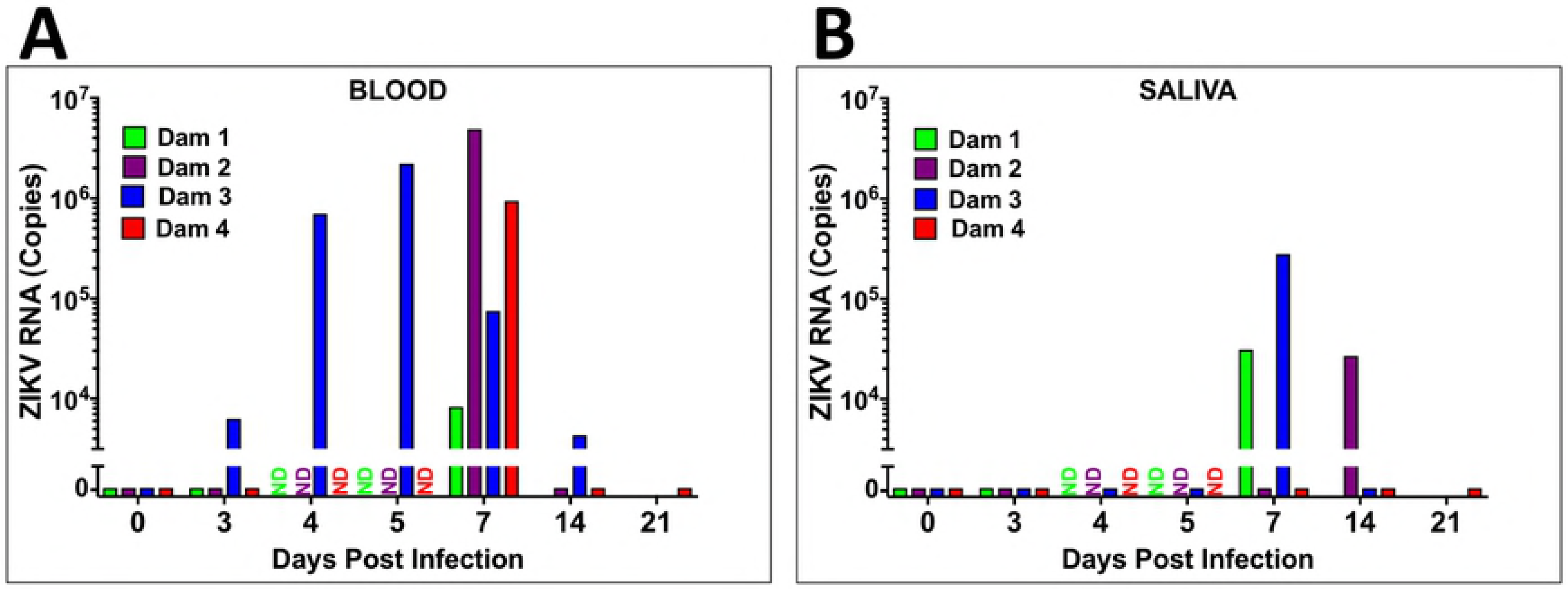
Viral loads (RNA) in plasma (A) and saliva (B) from ZIKV-infected pregnant baboons. A one-step qRT-PCR was used to measure ZIKV RNA in whole blood (A) and saliva (B) from each animal at indicated days post-infection (copies per milliliter; ND: not determined).

ZIKV RNA in urine was only detected in Dam 2 at 14 days post-infection (day of necropsy) (urine was not collected from Dam 3). Only Dam 4 had ZIKV RNA in CSF (day 7 post-infection). None of the dams had ZIKV RNA in vaginal swabs at any time point. Reproductive tissues (cervix, uterus, ovaries), cerebral cortex and cerebellum were examined for ZIKV RNA from the dams from tissues taken at the time of necropsy. ZIKV RNA was only detected in the uterus of Dam 2 (14 days post-infection).

- **Placental, amniotic fluid and fetal ZIKV Viral loads**

Fetuses are coded to match the dams (eg. Dam 1 = Fetus 1). We did not detect ZIKV in cord blood obtained at necropsy from any of the four fetuses. Fetus 2 (14 days post-infection) had ZIKV RNA in cerebral cortex, lung, spleen and ovary (Table 2). ZIKV RNA was found in Fetus 4 (21 days) in placenta, cerebral cortex, lung, spleen, intestine and ovary. ZIKV RNA was detected in amniotic fluid for both Fetus 2 and 4 (Table 1,2). Fetus 1 (7 days) and Fetus 3 (14 days) did not have detectable ZIKV in any tissue examined or amniotic fluid.

**Table 1.** Placental and amniotic fluid ZIKV RNA

| Baboon | Placenta |  |  | Amniotic Fluid |
| --- | --- | --- | --- | --- |
|  | Site 1 | Site 2 | Site 3 |  |
| Dam 1 | <LD | <LD | <LD | <LD |
| Dam 2 | $2.2 \times 10^6$ | $1.9 \times 10^4$ | <LD | $1.5 \times 10^5$ |
| Dam 3 | $1.5 \times 10^4$ | <LD | <LD | <LD |
| Dam 4 | $1.3 \times 10^4$ | $9.7 \times 10^4$ | $6.5 \times 10^4$ | $9.5 \times 10^5$ |
| (<LD: below limit of detection [$1 \times 10^2$ copies]; ND: not determined) | | | | |

**Table 2.**
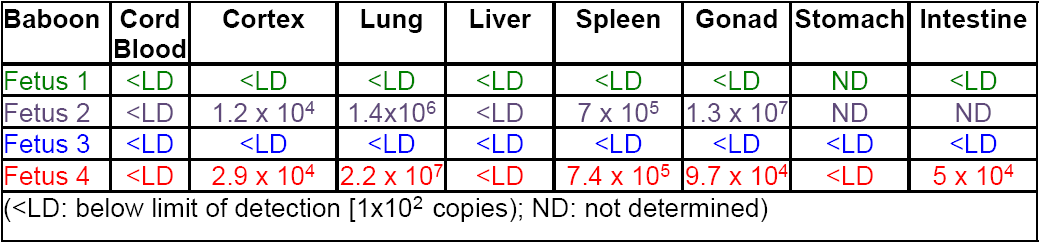
Fetal tissue ZIKV RNA

- **ZIKV specific IgM and IgG and neutralization (PRNT)**

**IgM**: All dams were negative for ZIKV IgM prior to infection and remained negative through 7 dpi. The three dams (Dams 2, 3 and 4) sampled at 14 days post-infection had IgM for ZIKV; Dams 3 and 4 exhibited a robust IgM response, Dam 2 had low ZIKV IgM, ∼25% that of the other dams. In Dam 4, ZIKV IgM remained constant from 14 through 21 dpi (Fig 3A).

**Figure 3.**
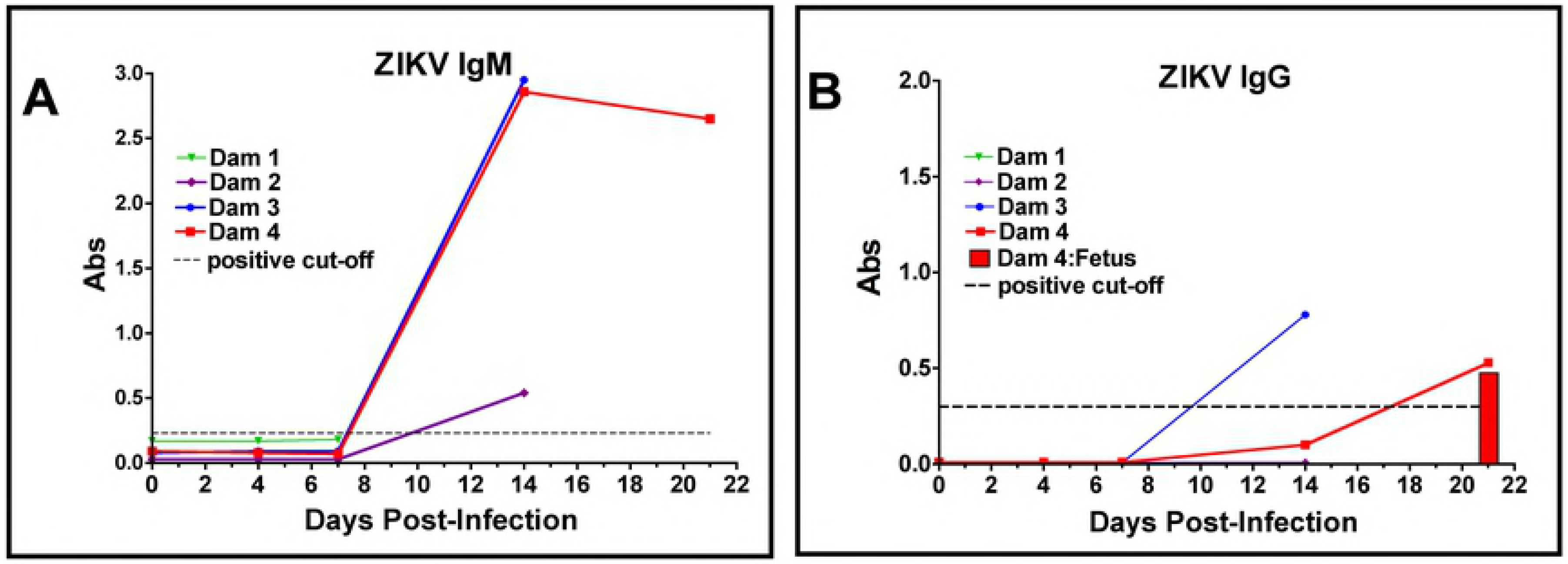
Detection of anti-ZIKV antibody responses in pregnant baboon serum. The presence of antibodies directed against ZIKV was determined by ELISA for IgM (A) or IgG (B). anti-ZIKV IgM were detected at 14 days post-infection in all three baboons sampled at this time point. Only two of the pregnant baboons had anti-ZIKV IgG; one at 14 days and the other by 21 days post-infection. The dam (2) with the weakest IgM response at 14 days did not exhibit an IgG response at this time point. The fetus of Dam 4 had anti-ZIKV IgG at levels similar to the mother indicating efficient FcγR transfer of IgG across the placenta.

**IgG**. No dams had serum ZIKV IgG by day sever post-infection. Dam 3 and 4 had ZIKV IgG by day 14; ZIKV IgG increased from 14 to 21 dpi in Dam 4. ZIKV IgG was also detected in the cord blood of Fetus 4 (14 dpi). Dam 2 did not have ZIKV IgG at 14 days; this dam also had low IgM (Fig 3B).

**PRNT.** In the two dams that exhibited an ZIKV IgG response, Dam 3 had a PRNT50 of 1:1280 (day 14) while Dam 4 had a PRNT50 of 1:2560 (day 21). The cord plasma from Fetus 4 had a PRNT of 1:640.

- **Plasma cytokine and chemokine response to ZIKV infection**

The cytokine/chemokine response was highly variable between dams. Dam 1 was notable in that no discernable change in any cytokine/chemokine was observed on day three or seven post-infection (study termination, Fig 4A). Dam 2 had an observable increase in IL-1β, IL-2, IL-6, IL-7, IL-12, IL-15, IL-16 and IL-17A, peaking on day 14 post-infection for all but IL-12 and IL-15 (day 7; Figure 4B). Dam 3 exhibited an increase in IL-2 (small), IL-6, IL-7, and IL-15; notably, all cytokines increased on day 7 and returned to basal by day 14 post-infection (Fig 4C). Dam 4 exhibited and increase in IL-1β, IL-2, IL-6, IL-7, IL-15 and IL-16 (Fig 4D), and similar to Dam 2, peak levels of these cytokines were on day 14 post-infection with the exception of IL-15 (day 7); by 21 days post-infection, most cytokines had returned to baseline or were lower than peak levels.

**Figure 4.**
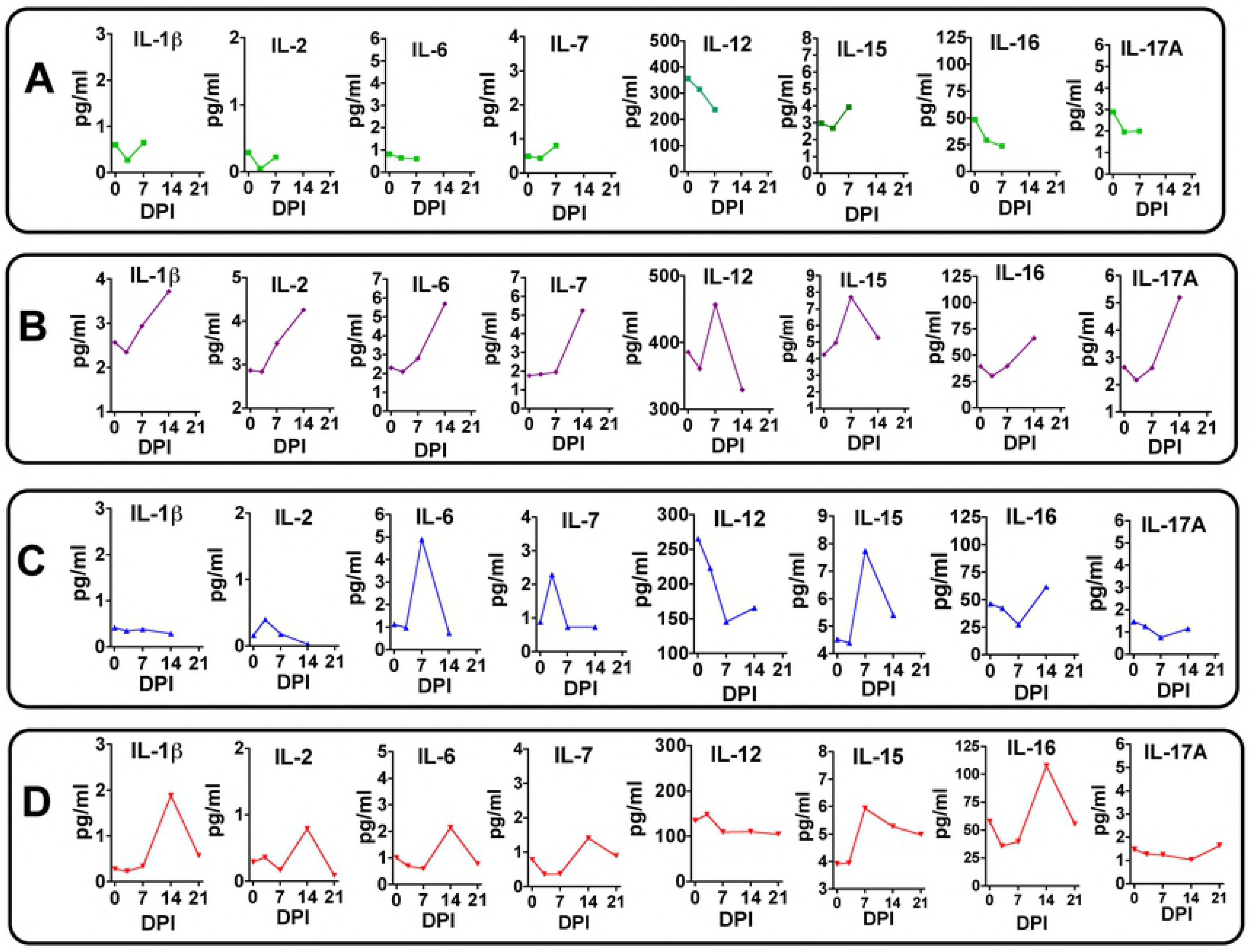
Maternal plasma cytokine concentrations in response to ZIKV infection in timed pregnant baboons. Only cytokines that were detectable and exhibited changes in concentration in response to ZIKV infection are shown. In Dam 1 (A) studied for 7 days post-infection, no chemokines increased post-infection. For Dam 2 (B), increases in IL-β, IL-2, IL-7, IL-12, IL-15, IL-16 and IL-17A were observed with peak levels either at day 7 or 14 post-infection. For Dam 3 (C) acute (day 7) increases were observed for IL-6, IL-7 and IL-15 (with a small increase at day 7 for IL-2). For Dam 4 (D) increases in IL-1β, IL-2, IL-6 IL-7, IL-15 and IL-16 were observed with peak levels either at day 14 (exception IL-15 at day 14), with resolution by study termination at day 21 for this dam.

Similar to that seen for cytokines, Dam 1 did not display any notable increase in plasma chemokine levels post-infection (Fig 5A). Dam 2 had notable increases in plasma levels of Eotaxin, MCP-1 and MCP-4 (Fig 5B), and similar to cytokines for this Dam, chemokines exhibited a progressive increase from Day 0 through study termination on Day 14. Dam 3 exhibited small transient increases in Eotaxin and IL-8 on day 7 (Fig 5C). Dam 4 had increases in plasma levels of Eotaxin (small), IL-8 and MCP-4 peaking primarily on day 14 post-infection (Fig 5D).

**Figure 5.**
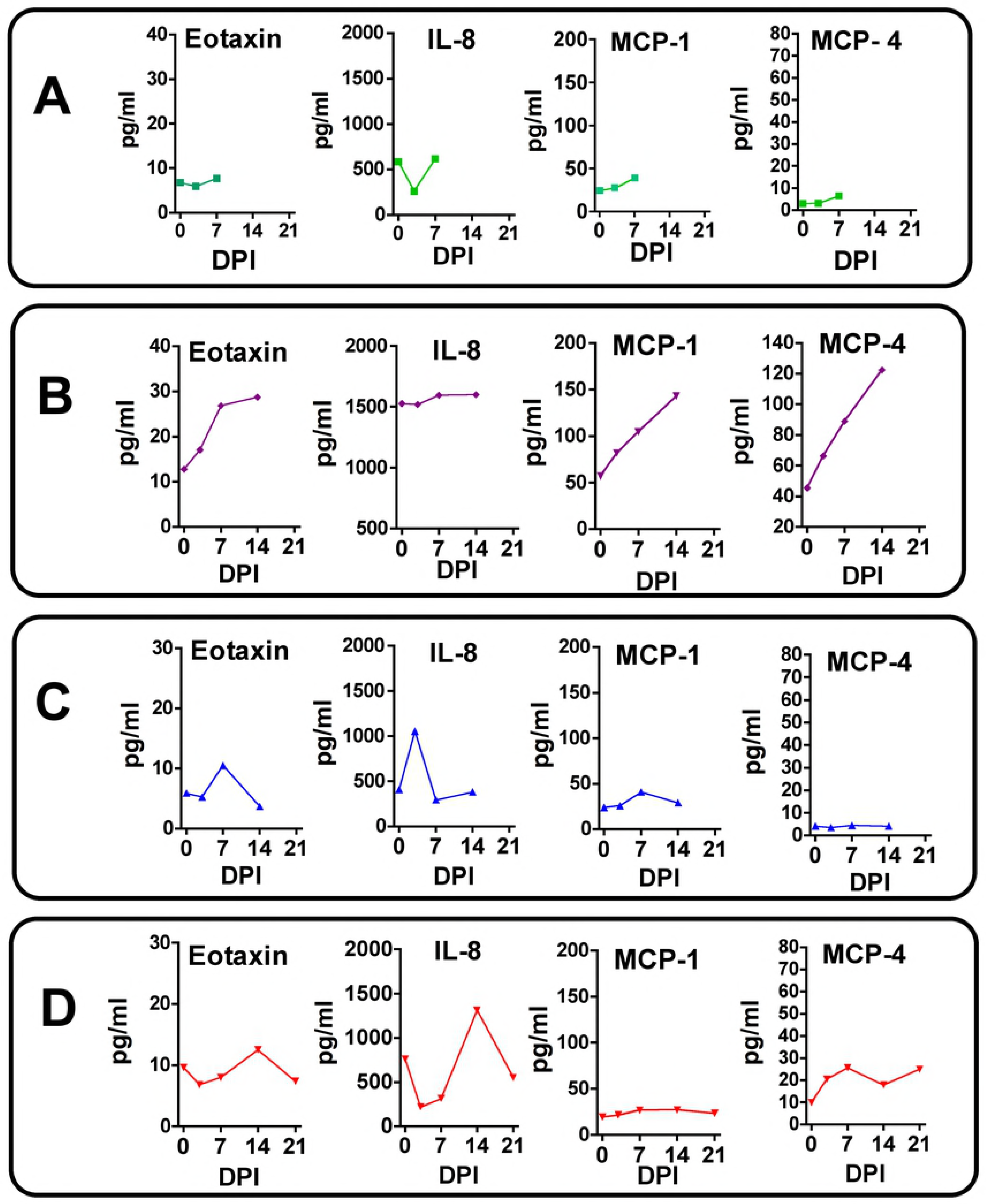
Maternal plasma chemokine concentrations in response to ZIKV infection in timed pregnant baboons. Only chemokines that were detectable and exhibited changes in concentration in response to ZIKV infection are shown. In Dam 1 (A) studied for 7 days post-infection, no chemokines increased post-ZIKV infection. For Dam 2 (B), increases in Eotaxin, MCP-1 and MCP-4 were observed peaking at day 14 post-infection. For Dam 3 (C) a transient increase in Eotaxin and IL-8 was observed. For Dam 4 (D), a similar transient increase in Eotaxin, IL-8 and MCP-4 was noted.

### CNS Immunohistochemistry

Upon standard H&E staining, none of the three fetuses with available CNS tissue for histology had demonstrable pathology of the cerebral cortex or other brain structures (Fetus 2 had extensive autolysis of the brain after *in utero* death). Histological examination of the frontal cortex (CNS region with abundant ZIKV RNA; 1×10^4^ copies/mg) of Fetus 4, which exhibited vertical transfer of virus at 21 days post-infection, revealed no major gross pathological lesions or decreased cortical volume compared to the control fetus or the two fetuses collected at days 7 and 14 post-infection with no evidence of vertical transfer of ZIKV.

- **GFAP (Glial Fibrillary Acidic Protein)**

Immunofluorescence (IF) for GFAP, a classical marker for radial glia (RG) and astrocytes in the developing cortex, revealed a pronounced difference in the ZIKV infected frontal cortex compared to the control (or day 7 or 14 post-infection fetuses without vertical transfer of virus). In the control fetus and fetuses with uninfected brains, the anticipated pattern of dense glial fibers projecting from the ventricular zone (VZ) to the marginal zone (MZ) was observed (Fig 6A, B, C, D). However, in the frontal cortex from the ZIKV infected fetus, there was a pronounced decrease in GFAP fibers, in particular in the subplate (SP) and intermediate zone (IZ; Fig 6E). Concurrent with the loss of RG fibers, a noted increase in the density in astrocytes was observed in the IZ/SP regions compared to the cortex of the control fetus and the two fetuses from infected mothers not exhibiting vertical ZIKV transfer. In these fetuses, GFAP-IF revealed a pattern of sporadic astrocytes with few astral branches representing maturing astrocytes with normal growing processes typical of this gestational age.

**Figure 6.**
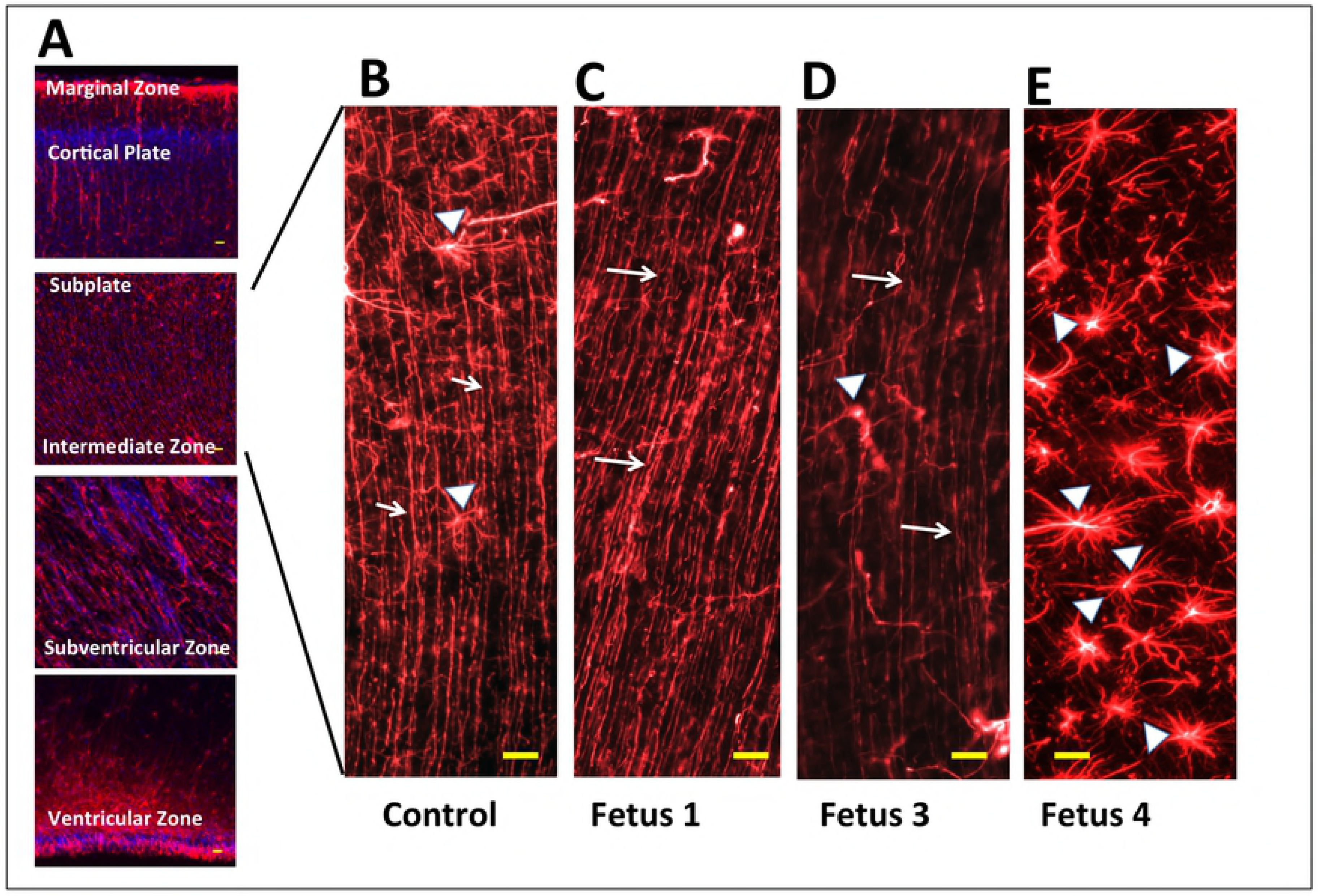
Immunofluorescence for GFAP in the frontal cortex of the fetal baboons. A reconstruction of GFAP-IF from marginal zone (MZ) and cortical plate (CP) through the subplate (SP), intermediate zone (IZ), subventricular zone (SVZ) and ventricular zone (VZ) in the control mid-gestation fetal frontal cortex (116 days gestation). B-E: GFAP-IF radial glial fibers (arrows) and occasional astrocytes (arrowheads) in the SP of the frontal cortex in control (B), and in three fetuses after maternal infection with ZIKV (C-E). Fetus 1 (C; 7 days post infection; 109 days gestation) and Fetus 3 (D; 14 days post infection;115 days gestation) had the anticipated pattern of GFAP-IF radial glia fibers comparable to the control fetus. Fetus 4 (E; 21 days post infection; 118 days gestation) that had detectable ZIKV RNA in fetal frontal cortex had virtually no continuous radial glial fibers, replaced with a dense population of astrocytes (arrowheads). (bar =20 μm).

- **Nestin and NeuN**

In order to determine if ZIKV infection targeted NPCs or reduced cortical neurons we performed IF for Nestin (NPCs) and NeuN (neurons, differentiating neurons). Immunofluorescence for Nestin was highly reduced in the frontal cortex of FM4 compared to control, in particular in the IZ/SP (Fig 7). When observed in the 21 day ZIKV positive cortex, Nestin IF cells were typically clustered, however there were large regions in the IZ/SP that were devoid of Nestin IF in the ZIKV infected fetus. In the control cortex, NeuN IF positive neurons were observed in long organized tracks of migration toward the CP, while in the ZIKV infected cortex, the pattern of NeuN IF appeared largely unorganized, although overall, the apparent number of NeuN neurons were similar between Fetus 4 and the control fetus (Fig 7).

**Figure 7.**
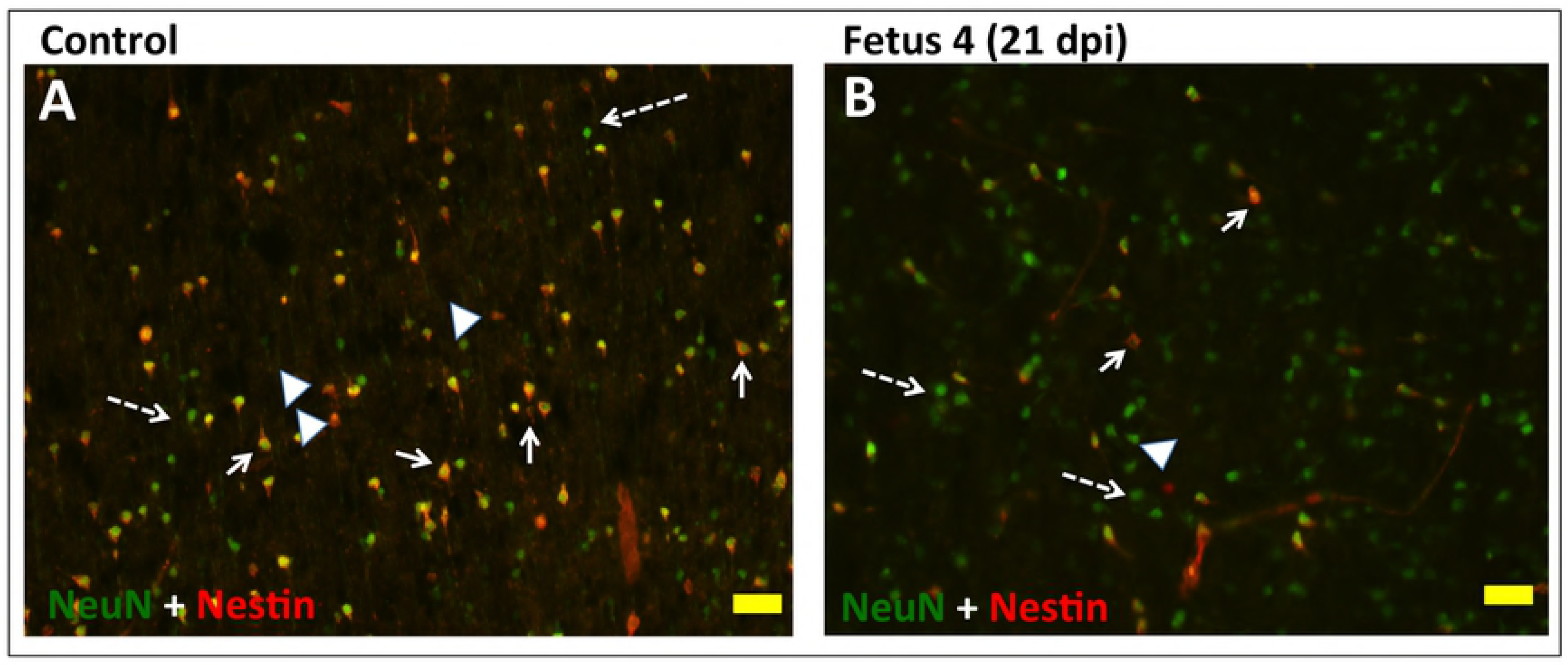
Double immunofluorescence for NeuN (green; differentiating neurons) and Nestin (red; neuroprogenitors) in the intermediate zone of the developing cortex of the control (A) and 21 day post ZIKV infection (B) fetal frontal cortex. There were a large number of cells co-expressing NeuN and Nestin in the control cortex (arrows; yellow) as well as NPCs (arrowheads) and differentiated neurons (dashed arrows). In the ZIKV infected Fetus 4 (B), there were similar densities of differentiating neurons/neurons (green) with only a few scattered remaining NPCs or cells co-expressing NeuN and Nestin. (bar=20 μm)

- **O1 (Immature Oligodendrocytes)**:

In order to determine if ZIKV infection causes white matter damage in term pregnancies and postnatally as reported in ZIKV^+^ fetuses/infants in human population (7) and observed in the 3^rd^ trimester pigtail macaque fetus with ZIKV positive brain (33), we performed IF for O1, a marker for immature oligodendrocytes and the only cell population that matures to oligodendrocytes that are responsible for myelinating the axons. O1 IF showed abundant O1^+^ immature oligodendrocytes in the subplate and IZ regions of the control fetal brain whereas fetus 4 (21dpi) brain showed decreased O1^+^ cells in both the subplate and IZ areas (Fig 8). O1 staining of cerebral cortices of Fetus 1 (7dpi) and Fetus 3 (14dpi) appeared similar to the control brain.

**Figure 8.**
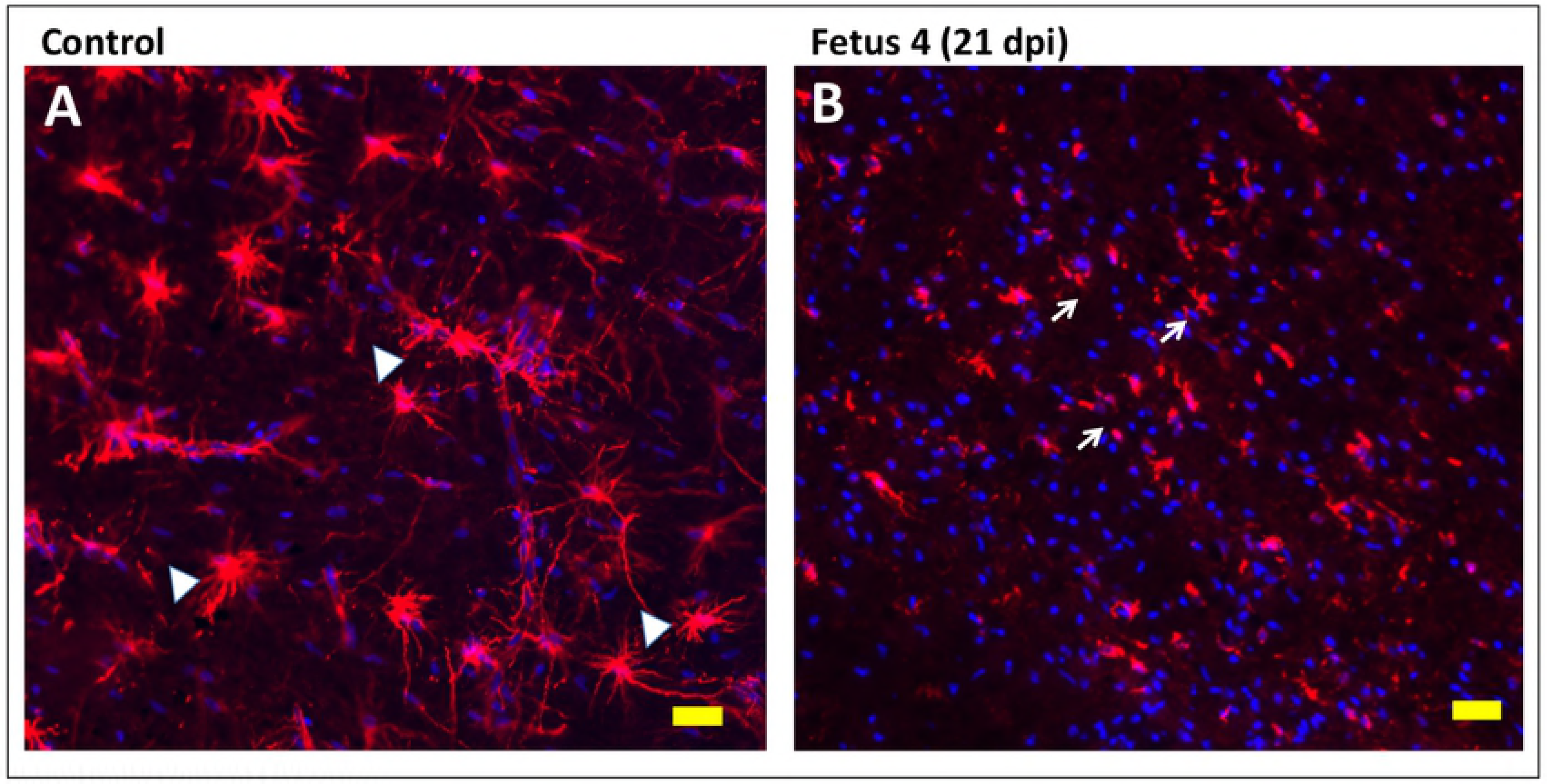
Immunoflurorescence for O1 (immature oligodendrocytes; OL) in the SVZ/IZ region of the developing fetal cortex. In the control fetal frontal cortex (A) there were numerous well developed immature OL (O1+; red) with extensive processes. In the ZIKV infected Fetus 4 (B), the number of O1+ cells were similar but morphologically distinct with limited processes and the appearance of degeneration or arrested development. (bar=20 μm; blue= DAPI stained nuclei)

- **Neuroinflammation**

The 21 day ZIKV infected cortex exhibited increased neuroinflammation with increased Iba1 (microglia) and IL-6 (proinflammatory cytokine) immunostaining (Fig 9D, H). Of interest however, was the noted increase in both Iba1 and IL-6 in the frontal cortex of the day 14 post-infection fetus despite not detecting ZIKV in the cortex of this fetus (Fig 9C, G). Neuroinflammation was not observed in the day seven post-infection frontal cortex that also did not have apparent vertical transfer of ZIKV at this time.

**Figure 9.**
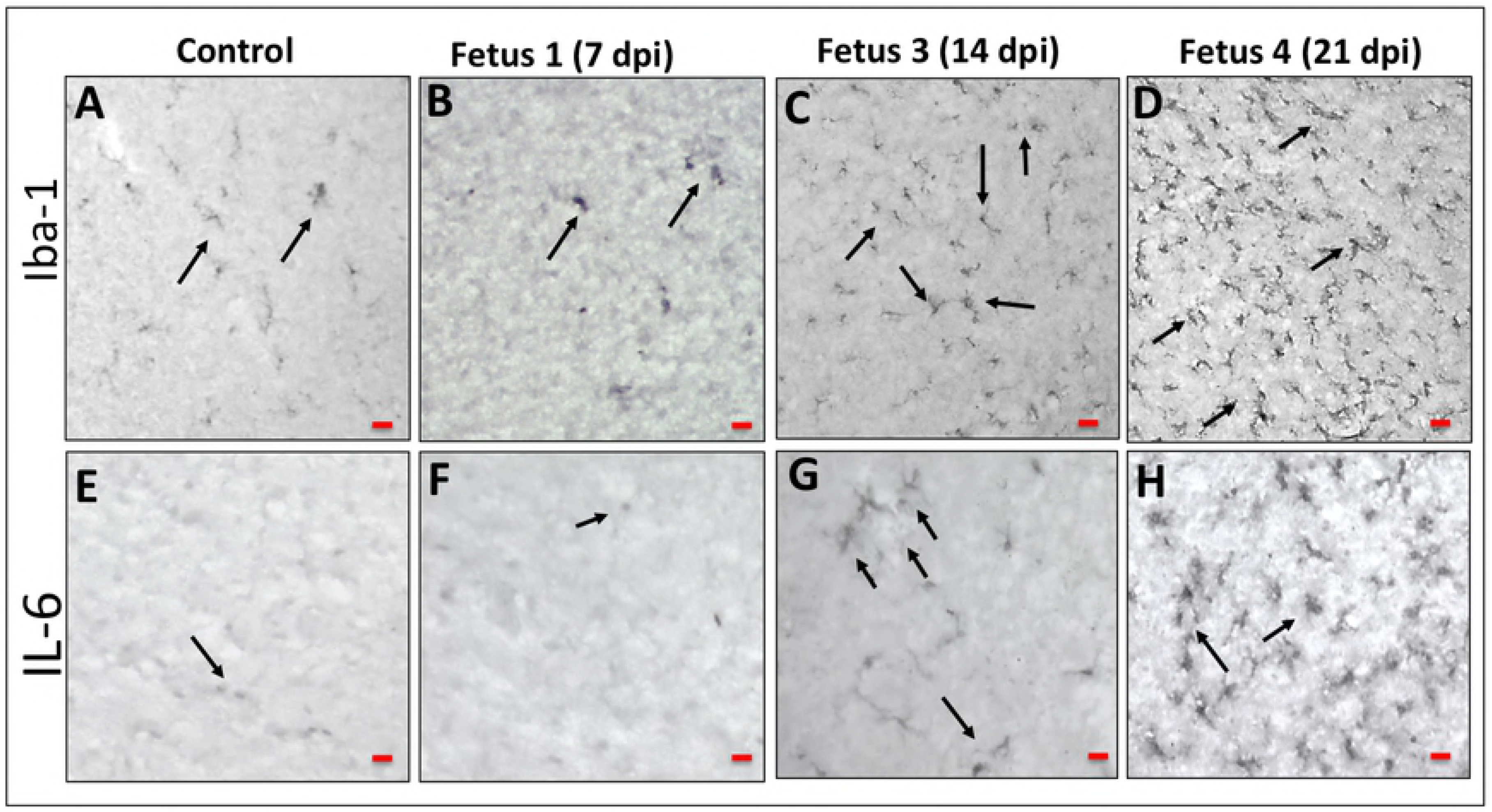
Immunohistochemistry for microglia (Iba1; A-D) and inflammation (interleukin 6 [IL-6]; E-H) in the frontal cortex of control (A,E) and fetuses whose mothers were infected with ZIKV. In both control (A) and Fetus 1 (7 days post infection; B), occasional Iba1 reactive microglia were observed (arrows). In the 14 day post infection fetus (C; Fetus 3) an increases number of Iba1 microglia were observed throughout the IZ through the CP. In the 21 day post infection Fetus 4 (D) this population increased to maximal. For IL-6 a similar pattern of immunostaining within the control and 7 day post infection fetus with only occasional random cells observed in the cortex (E, F). An increase in IL-6 staining was observed in the 14 day post infection fetus (G) with a pronounced IL-6 immunostained cells in Fetus 4 at 21 days post-infection.

- **Apoptosis (TUNEL)**

There were little to no apoptotic cells in the control frontal cortex in any region of the developing cortex (Fig 10A). There were notable apoptotic cells in the cortex in the day 21 post-infection cortex compared to the control cortex, primarily in the IZ/SP region (Fig 10B). The cortex of Fetus 1 was similar to the Control fetus with few apoptotic cells, while the day 14 post-infection fetus exhibited a similar amount of TUNEL staining compared to the ZIKV infected 21 day post-infection fetus.

**Figure 10.**
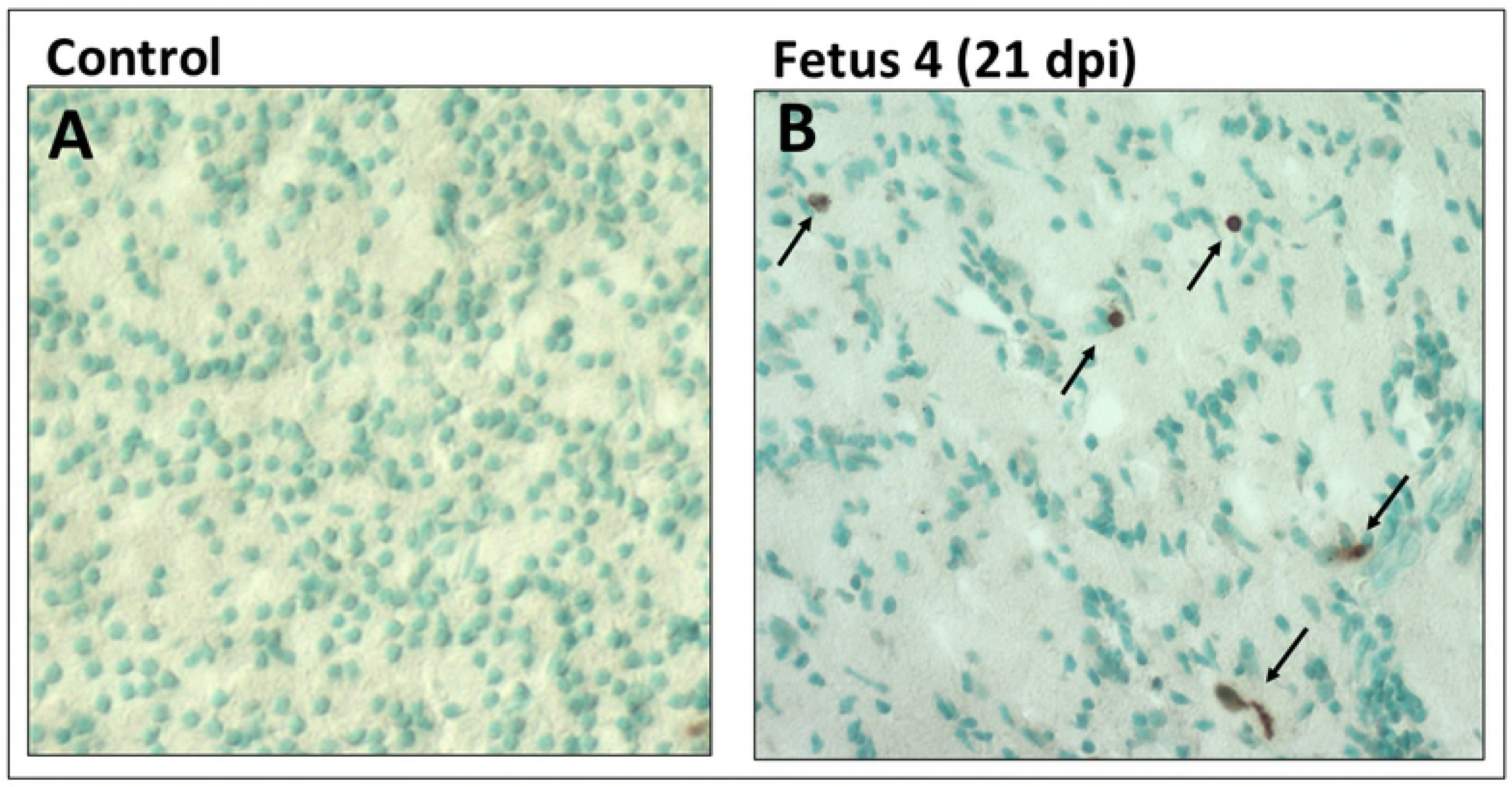
TUNEL staining for apoptosis in the frontal cortex of the control (A) and 21 day post-infection Fetus 4 (B). The cortex of the control fetus had very few apoptotic cells compared to the ZIKV infected fetus (brown cells; arrows). While there were dispersed TUNEL stained cells in the infected fetus, the density was not consistent with widespread apoptosis in the cortex of the infected fetus at 21 days post-infection.

## DISCUSSION

In this study we describe ZIKV infection of four olive baboons at near mid-gestation (97-107 dG; term ∼183 dG) following subcutaneous delivery of a relatively modest dose (1×10^4^ ffu) of the French Polynesian isolate resulting in vertical transfer of the virus to the fetus associated with fetal demise as well as significant fetal CNS pathology in two of the pregnancies. We chose the French Polynesian isolate since this strain was successful in achieving vertical transfer in pregnant rhesus macaques at the same dosage allowing for more accurate comparison of the outcome between the olive baboon and rhesus macaque (20, 35). In addition, a mutation in the prM protein (S139N) in the Asian strain of ZIKV that arose prior to the French Polynesian outbreak and has been stably maintained in the strains circulating in the Americas significantly enhances infectivity in human NPCs and yielding a more significant microcephaly in mice (41).

Following ZIKV infection, we observed that all four pregnant dams exhibited viremia within the first week post-infection and all presented with rash and conjunctivitis varying from mild to moderate. This differs from reports in macaques, male or non-pregnant and pregnant females where there have been few reports of rash and/or conjunctivitis. Similar to our findings in baboons, most clinical signs in pregnant humans, including rash, resolve within a week but may last up to two weeks. Description of rash following ZIKV infection in humans has been variable with estimates ranging from relatively infrequent (∼1/5^th^) (6) to greater than 2/3^rd^ of definite ZIKV cases in a Brazilian pregnancy cohort (45). As such, the presence and duration of rash and conjunctivitis in our pregnant baboons resembles that observed in human pregnancy compared to macaques.

While the magnitude of viremia achieved in the pregnant baboons was similar to that described in pregnant and non-pregnant macaques receiving a similar dose and route of delivery of ZIKV (French Polynesian or other strains), the onset of viremia in three of four pregnant baboons was delayed (detected at day 7 but not 3) compared to pregnant macaques which characteristically show sustained and prolonged viremia that initiates very early post-inoculation (1-2 days post-inoculation). Comparatively, viremia in these three pregnant baboons was cleared by day 14 post inoculation, whereas pregnant macaques typically display prolonged viremia with the viral RNA detectable in blood at one month or longer (20). However, a recent study of pregnant rhesus macaques did not find prolonged viremia despite using a higher inoculating dose (1×10^5^ pfu) of the Puerto Rican isolate (21). We did observe viremia prolonged to 14 days post-infection in one dam, and it is possible that this dam would have exhibited prolonged viremia if the time frame of the study had been extended. It is noteworthy that this dam also exhibited early viremia, detected at day three post-infection. In humans, viremia following ZIKV infection is usually short-lived (3-7 days) with occasional longer durations of up to 10 to 14 days (6). While prolonged viremia (46 to 53 days) has been reported in pregnant women(46), it should be noted that this was restricted to five cases after a search of the entire U.S. Zika Pregnancy Registry and as such, prolonged viremia during pregnancy in women appears to be rare. These authors did not find a correlation between prolonged viremia in women and an increased incidence of CZS, and as discussed below, vertical transfer of ZIKV was not observed in our dam with the longest duration of viremia.

Unlike the highly efficient vertical transfer of ZIKV described thus far in macaques (100%), we observed vertical transfer in two of four of the pregnant baboons infected at mid-gestation. In one dam, vertical transfer of ZIKV was associated with intrauterine fetal death by day 14 post-infection, while in a second dam, significant cerebral cortical neuropathology was observed in the fetus at day 21 days post-infection. The latter fetus was otherwise healthy with no congenital anomalies or signs of growth restriction. ZIKV RNA was detected in both fetuses in cerebral cortex, lung, spleen, and ovaries, and additionally in the intestine in the 21-day post-infection fetus. Unfortunately, considerable autolysis of the deceased fetal brain precluded meaningful histopathology in the case of intrauterine death. At the present time, we have not performed as comprehensive evaluation of all fetal tissues as has been done in macaque studies, instead, focusing in this report on major structures and the fetal CNS. ZIKV RNA was detected in the amniotic fluid and in the placenta of both of these pregnancies as well. We sampled three to four separate sites of each placenta representing different cotyledons since a recent study in macaques indicated that ZIKV infection of the placenta may be localized and not diffuse (31). In Dam 4, we found ZIKV RNA in three placental samples, while in Dam 2 (fetal death), we detected ZIKV RNA in two placental samples. We did not detect ZIKV RNA in the placenta of Dam 1 (study terminated at 7 days) but did in Dam 3 (one of three sites samples) despite not finding evidence of vertical transfer in this pregnancy. It can be argued that terminating study of Dam 1 at day seven post-infection may have precluded vertical transfer. Similarly, it is possible that vertical transfer of virus could have been delayed in Dam 3 as well, since we detected ZIKV RNA in the placenta and this dam also had the longest duration of viremia. Based on our observations, vertical transfer in baboons would appear to take place at some point between peak maternal viremia (7 dpi) and 21 dpi, and as such, a relatively early event during ZIKV infection. However, this does not preclude that vertical transfer could occur at a later time point post-infection since it has been suggested in women that vertical transfer may take up to 5 weeks. In support of our findings of a rapid transfer of virus to the fetus, Hirsch et al (31) observed vertical transfer in two late 2^nd^ trimester rhesus monkeys within 20 days of infection. In the other macaque studies it is unclear when vertical transfer occurred since pregnancies were studied for longer durations post-infection. While our limited study of four pregnant baboons would suggest that vertical transfer of ZIKV in baboons, unlike rhesus macaques, only occurs a subpopulation of infected pregnancies, we may have observed a greater frequency if we had extended the pregnancies in the two baboons terminated early. Clearly, future studies following infected pregnant baboons to longer periods post-infection are warranted. To our knowledge this is the first study to describe the early post-infection time course of vertical transfer of ZIKV in a primate.

Fetal death, miscarriage and preterm birth have been attributed to ZIKV infection in humans (5). There was a recent case report describing fetal death at 49-days post-infection in a rhesus macaque infected with the Puerto Rican strain (1×10^4^ pfu) of ZIKV during the 1^st^ trimester (30). In the other studies of pregnant macaques, ZIKV infection with either the Cambodian, French Polynesian or Puerto Rican isolates was not associated with fetal death or preterm birth despite allowing pregnancies to proceed until near-term gestation and with 100% vertical transmission of virus. In our case, fetal demise occurred just prior to two weeks post-infection. It is of interest that this dam exhibited a potentially delayed or suboptimal immune response to ZIKV with low IgM titers found at day 14 post-infection and an absence of IgG titers against ZIKV at day 14. This failure to mount an immune response in spite of the highest viremia may have contributed to the rapid vertical transfer and fetal demise. This dam also exhibited a notable systemic cytokine/chemokine response with progressive increases in plasma IL-1β, IL-2, IL-6, IL-7, IL-16, IL-17, MCP-1 and MCP-4 from day three through day 14, with acute increases in IL-12 and 15 that peaked at day 7. As such, these cytokines may have contributed to the adverse pregnancy outcome with fetal demise. Two other dams studied to 14 and 21 days displayed robust IgM titer on day 14 as well as having IgG titers and robust ZIKV neutralizing capacity by day 14 (Dam 3) and day 21 (Dam 4). Again, it is tempting to suggest that the earlier immune response in Dam 3 may have provided some protection to vertical transfer while the delayed IgG response in Dam 4 was insufficient to prevent vertical transfer even though there was efficient transfer of the IgG to the fetus. Unlike the case with fetal demise, Dam 3 exhibited a rather restricted cyto-chemokine response with elevation in plasma IL-6, IL-7, IL-8 IL-15 and IL-16 that resolved by day 14. However, more similar to Dam 2, Dam 4 had the delayed pattern of plasma cytokines (IL-1β, IL-6, IL-7, IL-8, IL-15 and IL-16) that peaked at day 14 post infection and returned to baseline by 21 days post-infection. Dam 1 was noteworthy in that there was no noted increase in plasma cytokines at either day 3 or 7 (study terminated) despite a robust viremia and rash.

Similar to that described in the rhesus, in our case of fetal demise (Dam 2), we noted rupture of fetal membranes at necropsy, consistent with the detection of ZIKV RNA in both amniotic fluid and urine. It is noteworthy that this is the only of the four infected dams in which we observed ZIKV RNA in urine, despite detecting the virus in the amniotic fluid of the 21-day post-infection dam (Dam 4) in which we also observed vertical transfer of ZIKV to the fetus. The fetal membranes of that dam were intact and the fetus otherwise appeared healthy with no meconium staining and normal weight for gestational age. Significant placental pathology has been described in macaques in response to ZIKV infection. Nguyen et al (35) reported minimal to moderate placentitis with variable calcifications in four of four pregnancies. Similarly, in a recent study by Hirsch et al (31) all five infected pregnancies exhibited at least microscopic placental infarctions with larger infarctions from earlier gestation infections as well as a subtle pattern of villous stromal calcification. Of interest, in the study of the single pigtail macaque infected with the high dose of Asian strain ZIKV, only mild deciduitis was noted which was also present in non-infected controls (33). In the present study, the only significant placental pathology was observed in the dam with intrauterine fetal death, exhibiting extensive fibrin deposition in the intervillous space with nearly uniform degenerated villi with frequent necrosis and acute inflammation. Two other placentas exhibited some minor evidence of inflammation (Dam 1 and 4) while the placental of Dam 3 was histologically similar to the control placenta, despite having one cotyledon positive for ZIKV RNA. It in this dam, it is possible that we did not histologically evaluate a placental region infected with the virus. Hirsch and colleagues observed that ZIKV infection in the placenta may be focal rather than diffuse with cotyledon by cotyledon variability in presence of ZIKV RNA in their study of rhesus macaques. We noted similar regional variation in ZIKV detection in placentas and as such, placental pathology may reflect restricted sampling of tissue for histology.

One of the major findings in the present study was a noted neuropathology in the 21 day post-infection fetal cortex with vertical transfer. Immunostaining with GFAP, a marker for Radial Glia (RG), revealed a pronounced difference in the ZIKV infected frontal cortex. Compared to the control fetus (and the two fetuses from ZIKV dams with no detectable vertical transfer) where we observed the anticipated pattern of dense radial glial fibers projecting from the ventricular to pial surfaces of the cortex, in the ZIKV infected frontal cortex there was a substantial loss of radial glial fibers, in particular in the IZ/SP region. In addition to serving as scaffolding for migrating neurons and neuronal precursors to the cortical plate to form the characteristic six-layered cortical structure via their fibers (47), RG also differentiate to neurons as well as astrocytes and pre-oligodendrocytes (48-51). Concurrent with the loss of the RG fibers, a dramatic increase in astrocytes was observed in the ZIKV infected frontal cortex. During normal cortical development in primates, a subset of GFAP-positive RG cease dividing in mid-gestation, presumably to serve as a scaffold for migrating neurons, then subsequently resume mitosis to generate astrocytes. Normal transformation of RG to astrocytes in the human fetal brain takes place gradually over several weeks from ∼15-35 weeks of gestation in the frontal cortex, occurring mainly in the second half of gestation (52). During this transformation, the RG end-feet detach first from the ventricular surface followed by release from the pial surface, while the RG cell bodies migrate upwards emerging in a bilaminar pattern with one band in the marginal zone/upper cortical plate and the other in the subplate/intermediate zones (SP/IZ) (53). Indeed, in the control fetus, GFAP staining revealed an astrocyte pattern reflective of this distribution, while in the ZIKV infected frontal cortex, a dense population of astrocytes was noted in the SP/IZ. This population of astrocytes in the ZIKV infected cortex suggests a rapid induction of differentiation of RG to astrocytes (and away from neuronal differentiation) rather than death of the RG *per se* concomitant with the noted loss of glial fibers. In the prior study in the pigtail macaque, vertical transfer of the Cambodian strain of ZIKV also resulted in an increase in GFAP-stained astrocytes in the white matter of the cortex as observed at six weeks after infection (at near-term gestation) (33). The findings of our study support that an early event following ZIKV penetration of the fetal CNS may be accelerated RG differentiation to astrocytes, in particular in the IZ/SP where the RG are primarily located during this stage of development. In mice, ZIKV delivery directly into fetal brains results in extensive microglial activation and astrogliosis, consistent with our findings(13). These authors noted that the GFAP immunostaining reflected a loss of RG and a progression of protoplasmic astrocytes into reactive astrocytes. Astrogliosis is a normal response to viral infection and brain injury and our results and that reported for the pigtail macaque are in agreement with this(54). In the study of the pigtail macaque (57), the authors suggested that ZIKV infection induced periventricular white matter injury resulting in the increased white matter gliosis and increased population of astrocytes. In humans, fetal RG have also been shown to differentiate into pre-oligodendrocytes (preOL), and since differentiation of preOL to immature/mature OL does not initiate until approx. 30 weeks of gestation in humans (∼138 dG in baboons)(55, 56), a loss of RG in the ZIKV infected cortex could conceivably reduce subsequent formation of OL and reduce myelination consistent with the observations in the pigtail macaque and as reported in human fetuses obtained from ZIKV infected pregnancies(33). O1^+^ immature cells of oligodendroglial lineage are the only cell population that initiates myelinogenesis by maturing into OL late in gestation when myelination of axons initiates (57). In the control fetus, we observed an abundant population of O1^+^ cells in a gradient from the IZ/SP through the CP of the control fetal cortex and developing white matter similar to what is observed in mid-gestation human fetus (58) exhibiting the characteristic multi-branched projections. In contrast, in the 21 day ZIKV infected fetal cortex, the O1+ cells were primarily without processes, fewer in number with the appearance of undergoing degeneration. In primates, cortical white matter forms within, and eventually replaces the IZ/SP and as such, the effects of ZIKV infection on both immature OL’s and astrocyte differentiation noted in our present study may disrupt normal white matter development (51). Again, these findings are consistent with that reported in the pigtail macaque in which a primary outcome from ZIKV infection later in the second trimester lead to a primary outcome of reduced white matter.

Studies in mice where ZIKV was infected directly into the fetal brains showed that a predominant outcome was loss of NPCs, either through apoptosis or altered cell-cycle regulation and decreased differentiation (13, 14). In our study, while TUNEL staining revealed a small increase in apoptosis in the 21 day post-ZIKV infection frontal cortex compared to the control, the degree of apoptosis does not appear to support a mass targeting of NPCs by ZIKV in the mid-gestation fetal baboon cortex. Alternatively, we cannot rule out that ZIKV-induced apoptosis occurred prior to 21 days post-infection, as an early event in ZIKV penetration of the fetal brain. We also stained for Nestin, a marker of NPCs including RG, as well as NeuN, a marker for differentiated neurons. In the ZIKV infected cortex, Nestin^+^ cells were highly reduced, in particular in the IZ/SP, and when observed were typically clustered. A loss in Nestin^+^ cells suggests that apoptosis of NPCs may have been an earlier event and that the outcome of ZIKV includes both a precocious differentiation of RG to astrocytes coupled with a loss of NPCs including RG due to apoptosis. In the control cortex, NeuN neurons were organized in long organized tracks of migration toward the CP, while in the ZIKV infected cortex, the pattern of NeuN staining was largely unorganized, although overall, the apparent number of NeuN neurons were similar. ZIKV infection in the baboon fetal cortex at mid-gestation does not appear to affect the already formed neuronal population in the cortical plate but could affect the final migration of neurons from the point of infection at mid-gestation onward since the SVZ in humans become the principal source of cortical neurons from 25 to 27 weeks of gestation. It should also be emphasized that the RG fibers provide an additional function in the formation of gyri and sulci (59). Loss of these fibers in the ZIKV brain would seemingly predict a less folded brain as gestation progresses. Loss of gyri/sulci is a hallmark in ZIKV infected cases of human microcephaly (8, 60). As such, we speculate that if allowed to mature to a later stage, the ZIKV infected fetus would likely have exhibited decreased brain folding as well as reduced cortical volume and white matter damage.

In addition to increased astrocytes, the 21 day ZIKV infected cortex exhibited additional indices of neuroinflammation with increased Iba1 (microglia) and IL-6 (proinflammatory cytokine) immunostaining. Of interest was the noted increase in both Iba1 and IL-6 in the frontal cortex of the day 14 post-infection fetus despite not detecting ZIKV in the cortex of this fetus. This implicates that the increased neuroinflammation may be in response to maternal or placental inflammation, or that ZIKV had transferred to the fetus in a tissue not sampled or in an adjacent region of the fetal brain not sampled. This dam did have a noted increase in plasma IL-6 at 7 days post-infection and this cytokine crosses the placental barrier. Increased neuroinflammation may also hallmark impending ZIKV infiltration into the fetal CNS. Neuroinflammation was not observed in the day seven post-infection frontal cortex that also did not have apparent vertical transfer of ZIKV at this time. The potential implications of increased fetal neuroinflammation in the absence of vertical transfer has implications for human infants from ZIKV infected mothers without notable gross CNS pathologies that may lead to subtle neurobehavioral or cognitive deficits post-birth. Indeed, delivery of the viral mimetic, poly (I:C) to pregnant macaques during the first or second trimesters results in offspring with notable behavioral changes reflecting autism (61).

In conclusion, the pregnant baboon offers an additional non-human primate model for ZIKV infection and adverse pregnancy outcome to compare and contrast with macaque species. Using a moderate dose of a relevant strain of ZIKV we recapitulated both clinical signs of human ZIKV infection as well as vertical transfer and a noted cortical pathology that may provide insight into the mechanisms via which ZIKV can induce fetal CNS damage in human pregnancy. Future studies can focus on long term neurodevelopmental sequelae resulting from ZIKV infection, in particular since we observed neuroinflammation of the fetal CNS in the apparent absence of vertical transfer of the virus. This may have major ramifications for the vast majority of cases in human pregnancies infected with ZIKV without CZS yet may have significant cognitive and behavioral deficits in the offspring.

## MATERIALS and METHODS

- **Ethical Statement**

All experiments utilizing baboons were performed in compliance with guidelines established by the Animal Welfare Act for housing and care of laboratory animals and conducted in accordance with and approval from the University of Oklahoma Health Sciences Center Institutional Animal Care and Use Committee (IACUC; protocol no. 101523-16-039-I). All studies with ZIKV infection were performed in Assessment and Accreditation of Laboratory Animal Care (AAALAC) International accredited ABSL2 containment facilities at the OUHSC. Baboons were fed standard monkey chow twice daily as well as receiving daily food supplements (fruits). Appropriate measures were utilized to reduce potential distress, pain and discomfort, including post-CSF collection analgesia. All animals received environmental enrichment. ZIKV infected animals were caged separately but within visual and auditory contact of other baboons to promote social behavior and alleviate stress. At the designated times post inoculation (Figure 1), the animals were euthanized according to the recommendations of the American Veterinary Medical Association (2013 panel on Euthanasia).

- **Animals**

Adult timed pregnant female olive baboons (n=5, 6-15 years of age) were utilized for this study. All females were multiparous with history of successful prior pregnancies.

- **Virus stocks, Infection and sample collection**- Animals were anaesthetized with an intramuscular dose of Ketamine (10 mg/kg) before all procedures (viral inoculation, blood, salivary and vaginal swabs and urine collection). Timed pregnant female baboons were infected subcutaneously at the mid-scapular area with a single clinically relevant dose of 10^4^ focus forming units (ffu; 1 ml volume per dose) of the French Polynesian ZIKV isolate (H/PF/2013). The dosage used to infect the animals in our study is based on the previous works done in mosquitoes carrying WNV and DENV, where it was estimated that mosquitoes carry 1×10^4^ to 1×10^6^ plaque forming units (PFU) of the virus (62), from a study evaluating Brazilian ZIKV in a bite from *Aedes aegypti* mosquito(63) and from a study of mosquito transmission of ZIKV in rhesus monkeys (34). The pregnant females were infected near mid-gestation (between 97 and 107 days of gestation [dG]; term is approx. 181 dG; the overall approach is detailed in Fig 1). Maternal blood samples, vaginal and salivary swabs and urine was obtained on the day of inoculation (day 0) as well as ultrasound evaluation of fetal viability. Whole blood was collected into EDTA tubes. Urine was collected by direct bladder cystocentesis. Saliva and vaginal samples were collected by cotton roll salivette. The sampling procedure for each dam is detailed in Fig 1. For Dam 1, blood, urine, vaginal, salivary and CSF samples were obtained at 3 dpi and at 7 dpi, immediately prior to euthanasia at 7 dpi and maternal-fetal tissue collection. For Dam 2, blood, urine, vaginal, salivary and CSF samples were collected at 3 dpi and 7 dpi and at 14 dpi, immediately prior to euthanasia (14 dpi) and maternal-fetal tissue collection. For Dam 3 blood, vaginal, salivary samples were collected on days 3,4,5 and 7 dpi and at 14 dpi, immediately prior to euthanasia (14 dpi) and maternal-fetal tissue collection (no urine or CSF collection for this Dam). For Dam 4, blood, urine, vaginal, salivary and CSF samples were collected at 3, 7and 14 dpi, and at 21 dpi, immediately prior to euthanasia (21 dpi) and maternal-fetal tissue collection. A control timed pregnant dam was euthanized on 116 dG.

At the end of the study for each animal, dams were sedated with ketamine, all maternal samples obtained as well as ultrasound measurements, then the animal rapidly euthanized with euthasol. A C-section was quickly performed, cord blood obtained and the fetus euthanized with euthasol. Maternal and fetal tissues were rapidly collected and samples were both fixed with 4 % paraformaldehyde and frozen on dry ice (stored at −80°C) for each tissue.

- **One-Step quantitative reverse transcription PCR** – Primers and probes used for qRT-PCR were designed by Lanciotti et al (64) (**Table 3**). RNA was isolated from maternal and fetal tissues (**Table 1**) using QIAamp cador pathogen mini kit (Qiagen, Valencia, CA). ZIKV RNA was quantitated by one-step quantitative real time reverse transcription PCR using QuantiTect probe RT-PCR kit (Qiagen) on an iCycler instrument (BioRad). Primers and probes were used at a concentration of 0.4 μM and 0.2 μM respectively and cycling conditions used were 50°C for 30 min, 95°C for 15 min followed by 40 cycles of 94°C for 15 s and 60°C for 1 min. Concentration of the viral RNA (copies/milliliter) was determined by interpolation onto a standard curve of six 10-fold serial dilutions (10^6^ to 10^1^ copies/ml)) of a synthetic ZIKV RNA fragment available commercially from ATCC (ATCC VR-3252SD). The cutoff for limit of detection of ZIKV RNA was 1×10^2^.

**Table 3.**
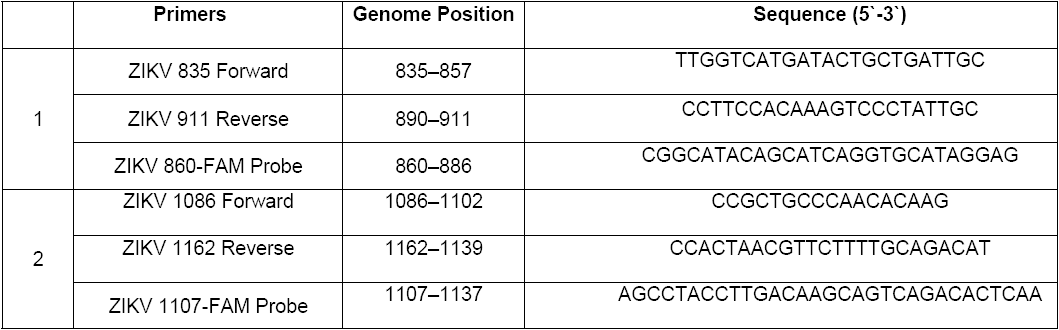
Primer/Probe sets for the detection of ZIKV by one step qRT-PCR

- **ZIKV ELISA** - ZIKA specific IgM and IgG antibody responses were assessed in the serum samples using the commercially available anti-ZIKV IgM (#ab213327, Abcam, Cambridge, MA) and IgG (#Sp856C, XpressBio, Fredrick, MD) ELISA kits. Briefly, a 1:100 for IgM and 1:50 for IgG serum dilution was performed in duplicate and added to the precoated plates available in the kits. The assays were performed using the manufacturer’s instructions and the assay was read at 450 nm for IgM and 405 nm for IgG antibodies in the serum.
- **Plaque Reduction Neutralization Test (PRNT) -** A plaque reduction neutralization test (PRNT) was used to assess serum samples for ZIKV neutralizing antibodies. Vero cells (ATCC #CCL-81) were maintained in DMEM (supplemented with 10% heat-inactivated FBS, 1x antibiotic/antimycotic), seeded in 12-well plates (2 ×; 10^5^ cells/well, 37° C, 5% CO_2_) and incubated at for approximately 24 hours until 80-100% confluent. Baboon sera were heat inactivated (56° C, 30 min) then serially diluted 2-fold in media, followed by addition of 200 plaque forming units (pfu) of French Polynesian ZIKV isolate (H/PF/2013). Samples were vortexed and incubated in a 37° C water bath for one hour. Media was then removed from each well and replaced with the virus/serum mixture followed by incubation at 37° C for one hour with intermittent rocking of the plates every 20 minutes. Control wells included inoculation with 200 pfu of ZIKV with no serum added to determine total plaques, as well as control wells without virus. A 1% carboxymethyl cellulose overlay was then added without removing the inoculum and the plates were incubated at 37° C for approximately 72 hours. The cells were then fixed with 1% paraformaldehyde for one hour, after which, the overlay was removed and the cells were stained with 1% crystal violet. The PRNT50, or the concentration of serum required to neutralize 50% of the plaque count of a known amount of serum-free ZIKV, was calculated by visually counting plaques.
- **Serum Cytokine Analysis** – Non-human primate cytokine/chemokine/inflammatory panel V-PLEX Multi-Spot assay system (Meso Scale Discovery, Rockville, MD) was used to quantify 24 cytokines and chemokines from plasma obtained from dams and cord blood. The assays were performed according to the manufacturer’s instructions. Briefly, plasma samples were diluted 2-fold (inflammatory, cytokine panel) and 4-fold (chemokine panel) in respective diluents and added in duplicates to plates for each panel and incubated overnight on a shaker at 4°C. Plates were washed 3 times and detection antibody cocktail specific for each panel was added to the respective plates and incubated for 2 hours at room temperature on a shaker. Plates were washed and read using an MSD instrument and the final data was obtained using the MSD discovery workbench software.
- **Fetal CNS Immunohistochemistry/Immunofluorescence –** Following removal, fetal brains were divided mid-sagittal with one half rapidly frozen on dry ice and stored at −80°C and the other half fixed in 4% paraformaldehyde for 24 hours, cryoprotected in 30% sucrose until sunk and frozen in embedding molds in optimal cutting temperature (OCT, Sakura Finetek) on dry ice and stored at −80°C. Serial coronal sections (30um) were cut on a freezing microtome and collected in 24 well plates in a cryoprotectant solution at −20°C. For Iba-1 and IL-6 immunohistochemistry, every fifth free-floating section was selected for immunolabeling. Briefly, sections were treated with 2% H_2_O_2_ solution to inactivate endogenous peroxidase activity followed by blocking and primary antibody incubation overnight at room temperature with rocking (Iba-1; Rabbit polyclonal 1:100, NBP2-16901; IL-6; Rabbit polyclonal 1:200, NB600-1131; NovusBio, Littleton, CO). Sections were washed and incubated with goat anti-rabbit HRP conjugated secondary antibody for 1 h at RT. Immunolabeling was visualized using DAB substrate and sections were washed and dried overnight before a cover slip was added with Permount (FisherScientific, Fairlawn, NJ). Control reactions were performed where the primary antibody was omitted from the procedure. Sections were visualized using a brightfield microscope (Olympus B40x) equipped with a SPOT 5 MP digital camera with SPOT 5.3 imaging software (Sterling Heights, MI).

For single and dual Immunofluorescence labeling (IF), sections were treated with 1% NaBH4 solution to reduce auto-fluorescence then blocked for an hour in blocking solution followed by overnight incubation in single or double primary antibodies (GFAP; mouse monoclonal 1:1000, NBP1-05197; Nestin; chicken polyclonal 1:1000, NB100-1604, Novus Bio, Littleton, CO; NeuN, mouse monoclonal 1:100, MAB377, Millipore, Temecula, CA; O1, mouse monoclonal 1:20, MAB1327, Bio-Techne, Minneapolis, MN). Sections were incubated for an hour the next day at room temperature in goat anti-mouse Alexa 568 and goat anti-chicken Alexa 488 1:200 (Life Technologies, Carlsbad, CA). Sections were washed in PBS and transferred to slides, incubated for 30 min in dark at room temperature in Prolong Gold with DAPI (Life technologies, Carlsbad, CA), cover slipped and cured for 24 h before visualizing using fluorescent microscope (Zeiss Axiovert 200 inverted fluorescent microscope). Controls again were exclusion of the primary antibodies. Images were captured using AxioVision imaging software (Zeiss, Germany).

- **TUNEL Assay (Apoptosis) –** Terminal deoxynucleotidyl transferase dUTP nick end labeling (TUNEL) assay was performed using a commercially available kit from Trevigen (Gaithersburg, MD). Briefly, fixed frozen fetal cortical sections were washed with PBS and incubated with Cytonin for 30 min, washed, and quenched before labeling with biotin-labeled dUTP. The labeling reaction was stopped by adding stop buffer. The tissue sections were then incubated with HRP-conjugated streptavidin for 10 min, washed, and immersed in DAB solution for color development. Sections were counterstained with methyl green before mounting. A positive and a negative control sections were included in the protocol. Apoptotic cells are stained with dark brown color.

## Acknowledgements

Oklahoma Baboon Research Resource for providing baboons; James Tomasek, Vice President for Research, University of Oklahoma Health Sciences Center, for support of the project; and Helen Lazear, University of North Carolina Chapel Hill, for providing ZIKV stocks.

